# PulseNet international: the missing link between PFGE and WGS

**DOI:** 10.1101/2021.03.08.434411

**Authors:** Ibrahim-Elkhalil M. Adam

## Abstract

**Introduction:** DNA-based surveillance of bacterial diseases has been using pulsed field gel electrophoresis (PFGE) since 1996. Currently, the international surveillance network (PulseNet international) is turning toward whole genome sequencing (WGS). ATCGs of WGS are compared using several sequence alignment methods. While patterns of horizontal lines of PFGE profiles are being compared using relative positioning of bands within a range of tolerance. A recently suggested image analysis algorithm and a deployed database (geltowgs.uofk.edu) collectively invented a promising method for comparing PFGE to *in-silico* obtained digestion models (DMs) derived from WGS. The database requires a parameter that determines PFGE resolution. Here, the author suggests a new method for calculating this factor. Epidemiological and molecular conclusions returned by the database are evaluated.

**Methodology:** two PFGE profiles representing *Xba*I digests of *E. coli* and *Salmonella enterica* analyzed by the suggested image analysis algorithm were submitted to the database after calculating resolution of PFGE using Dice percentage of difference between the closest PFGE bands in length. *E. coli* and *Salmonella enterica* test subjects were compared to 489 and 401 DMs respectively. The three data sets returned were analyzed.

**Results and conclusions:** according to modified PFGE evaluation criteria; a single DM is possibly related to *E. coli* test subject. It belonged to the same serovar. No epidemiologically related DM was shown for *S. enterica* test subject. Conclusions mentioned earlier could never be made ignoring co-migration. Standardization of both; suggested image analysis and database algorithms will deepen our understanding of bacterial epidemiology by means of possible qualitative approach built upon identification of fragment sequences and their locations within chromosomes.

## Introduction

In order to reveal genomic fingerprints of pathogenic bacteria in cases of suspected outbreaks, DNA digests are obtained using restriction enzymes with rare recognition sequence across chromosomes of interest. Different digestion protocols (restriction enzymes and running conditions) have been standardized for several bacterial species. DNA fragments are separated using pulsed-field gel electrophoresis (PFGE) [1]. This technology was initiated and being used for this purpose in USA since 1996 [2]. An international consortium of laboratories is growing ever since, building PulseNet international (PNI) [3,4]. Currently, 83 countries around the world have already joined PNI [5]. In case of PFGE typing, other phenotypic characteristics such as virulence associated mutations and antibiotic resistance profiles of isolates must be investigated using different methods [6]. Due to recent sophistication in DNA sequencing technology, PNI started to shift from PFGE to whole genome sequencing (WGS) of bacterial isolates [4]. Identification of strains, valuable information about their antibiotic resistance capabilities and of course genetic relationships to other isolates of previous outbreaks are provided by WGS [7,8].

In case of shifting to WGS without a method that links PFGE data accumulated during the last 26 years to WGS results, valuable information about bacterial population genetics, population dynamics and evolution will be lost. Taking into account that > 800,000 PFGE record are available in PulseNet network of USA alone [2]. Shifting to WGS will take time for all participant laboratories across 83 countries to upgrade their infrastructure for WGS [9]. Consequently, valuable epidemiological information will be also lost.

The simple logic behind PFGE is that number and distribution of recognition sequence of the restriction enzyme used determine the final profile [10]. While whole chromosome and plasmid DNA sequences revealed by WGS can also be digested *in-silico*, resulting in digestion models (DMs). It may seem possible to link PFGE to WGS data. A recently published scientific article [11] identified the following obstacles for doing so;

1. DNA fragments those have the same or very close molecular weight (up to 5% [12]) will resolve in a single PFGE band (fragments co-migration) [13].
2. Length of PFGE bands are estimated based on a DNA ladder’s exponential correlation between retention factors (rF) and band sizes [14].
3. Fragment length of *in-silico* digested WGS are exact numeric values, not simply estimations like the case in PFGE.
4. Typable PFGE fragments are limited by largest and smallest band sizes of the DNA ladder used [10].

Authors of the same article suggested an image analysis algorithm that quantitatively reveal number of co-migrated fragments across PFGE profiles. They named the new parameter “factor of co-migration (FCM)” [11]. In order to assess reliability of FCM calculations, they developed “WGSToGel”. Which are simulation algorithms that predict PFGE profiles in terms of expected band sizes and corresponding FCMs from *in-silico* digestions of whole chromosome sequences. They deployed geltowgs.uofk.edu; a freely accessible online database. It compares PFGE-FCM image analysis results to *in-silico* digested whole chromosome sequences [11].

In this study, the author suggests a new method for estimating the parameter critical co-migration threshold (CCT) required by GelToWGS database. Test subjects are previously published PFGE-FCM results of two widely used PFGE markers:

1. *Xba*I digestion profile of *E. coli* O157:H7 strain G5244 [13].
2. *Xba*I digestion of *S. enterica* serotype Braenderup (strain H9812) suggested by Hunter and her team [11].

Epidemiological conclusions returned by the database were analyzed. Detailed descriptions of their suggested methods are cited [11]. The author will also discuss a qualitative approach to analyze GelToWGS results. Considering DNA-based perspective of fragment location within the chromosome and epidemiological metadata retrieved from NCBI-BioSample database.

## Methodology

### Test subjects

The two PFGE markers were chosen for this study because they are widely used across PNI laboratories [14]. Factor of co-migration using exponential correlation between single-fragment bands and pixel densities (FCM-ECSB) is the image analysis algorithm suggested by Adam IE and his colleagues [11]. It depends on the observation that; across a PFGE profile, pixel densities (PDs) reduces as band size do [15]. Warner and Onderdonk reported that higher PDs shown by smaller band sizes are caused by co-migration [16]. FCM-ECSB simply calculate FCMs by calculating hypothetical values for bands with “odd PDs” based on other bands of the profile [11]. Other type of PFGE contamination (known as “ghost bands”) result in low PDs [17].

FCM-ECSB results of *E. coli* strain G5244 were reported in Supplementary material 1 (S1) of the cited article [11]. It is a screen-recorded video available online here. FCM-ECSB results of *S. enterica* strain H9812 were also reported by the same author and his colleagues [11].

### Selecting CCT and marker errors

The author suggests selection of the lowest value of Dice %age of difference [11] (between the closest band sizes across a given PFGE profile) as the CCT. Sine running distance is a crucial factor that determines resolution of PFGE [13].

Accordingly; for *S. enterica* test CCT was calculated between PFGE bands #10 (173.4 kbp) and 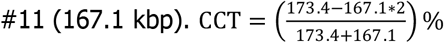 This results in 3.7. Similarly, CCT for *E. coli* test calculated between bands #19 (40.5 kbp) and #20 (36.3 kbp) resulted in 5.4687. PFGE marker errors for both test subjects as reported in mentioned citation [11] i.e., 0.007 and 0.002 for *S. enterica* and *E. coli* respectively.

### GelToWGS database

FCM-ECSB result for each test was uploaded in MS Excel workbook (.xlsx file is required). Corresponding parameters (digestion models, CCT and marker errors) were supplied to the system. GelToWGS database server returns three data sets for each test; The first shows over all summary of matches with each PFGE band. The second shows matches with each DM those share ≥ 5 DNA fragments alongside their metadata. Third data set represents details of matched and none matching fragments for each DM. These data sets are available with this publication in supplementary data sets 1 and 2 for *E. coli* and *S. enterica* tests respectively.

## Results

At selected CCTs for both test subjects, DMs those showed at least 5 matches with each test were 50.1% and 40.65% for *E. coli* and *S. enterica* respectively. *Xba*I digestions of *E. coli* chromosomes result in significantly higher number of typable DNA fragments than *S. enterica* (7,465 compared to 3,020).

Isolate relationships were critically affected by co-migration. For both tests, raw number of query matches was higher than typable matches. This will make no sense if FCM of each PFGE band is unknown. But when taking into account FCMs of test subject, number of actual matches was reasonable **Table (1)**.

**Table (1):** PFGE-FCM results and corresponding parameters submitted to GelToWGS database and summary of results returned by the database.

| Test data submitted to GelToWGS database |  |  |  | Compared to | Summary of results returned by GelToWGS database server |  |  |  |  |  |
| --- | --- | --- | --- | --- | --- | --- | --- | --- | --- | --- |
| Test subject | Total number of DNA fragments (PFGE-FCM) | Critical co-migration thresholds | PFGE marker error |  | Models having ≥ 5 matches |  |  | Matches with test subjects |  |  |
|  |  |  |  |  | No. of *In-silico* digested chromosomes | %age From total | typable fragments | Total query matches | Excluded due to co-migrations | Actual matches |
| *XbaI* digestion profile of *E. coli* O157:H7 strain G5244 | 25 | 5.4687 % | 0.007 | 489<br>*XbaI* –digested chromosome sequences of *E. coli* | 245 | 50.1% | 7,465 | 9,721 | 3,423 | 6,298 |
| *XbaI* digestion of *S. enterica* serotype Braenderup (strain H9812) | 20 | 3.7 % | 0.002 | 401<br>*XbaI* –digested chromosome sequences of *S. enterica* | 163 | 40.65% | 3,020 | 3,542 | 1,296 | 2,246 |

Significant numbers of matches were returned by GelToWGS database for both test subjects. But according to PFGE comparison criteria [18], a single DM in *E. coli* test was possibly related (differed by 6 fragments). Results of *S. enterica* test did not show any significant relationship according to the same criteria [18]. Details of *E. coli* and *S. enterica* results are in Supplementary data sets 1 and 2) respectively **Table (1)** summarizes findings.

### GelToWGS comparisons with *Xba*I digestion profile of *E. coli* O157:H7 strain G5244

Out of total query matches, 3,423 were excluded due to co-migration. Consequently, actual matches reduced to 6,298. These results suggest that DMs of *E. coli* stored in the database shows a high degree of overall similarity and 15.63% of matches mislead PFGE evaluation **Table (1)**.

In fact, top 26 DMs (with the exception of two clones) those have shown high similarity with test subject belongs to the same serotype (O157:H7). Among this number, sum of actual matches was 812 (12.89% of total matches). Number of these typable DNA fragment per each DM ranged from 23-34. Number of matched fragments with test data (19 PFGE band representing 25 DNA fragments) ranged from 19-14 within the O157:H7 serotype. According to currently adopted criteria [18,19], a single DM is considered possibly related to test subject; It is the strain 3384 with NCBI-GeneBank accession number CP017440.1 which showed Dice coefficient of 0.667, Supplementary data set 1 (S1).

Epidemiological information as retrieved from NCBI-BioSample database (Key: SAMN04447623) show that it was collected in USA Kansas state in 2003. It showed enterohemorrhagic activity (EHEC).

When considering possible occurrences of each PFGE band across DMs; band #6 (244.1 kbp, 1 DNA fragment) occurs in 419 out of 489 DM. query matches with this fragment ranged from 0 to 4. But when considering actual matched; band #5 (276.9 kbp representing two fragments) occurs the most. PFGE band #1 is very rare. It occurred in 48 and 50 times as actual and query match respectively **Fig. (1)**.

**Fig. (1):**
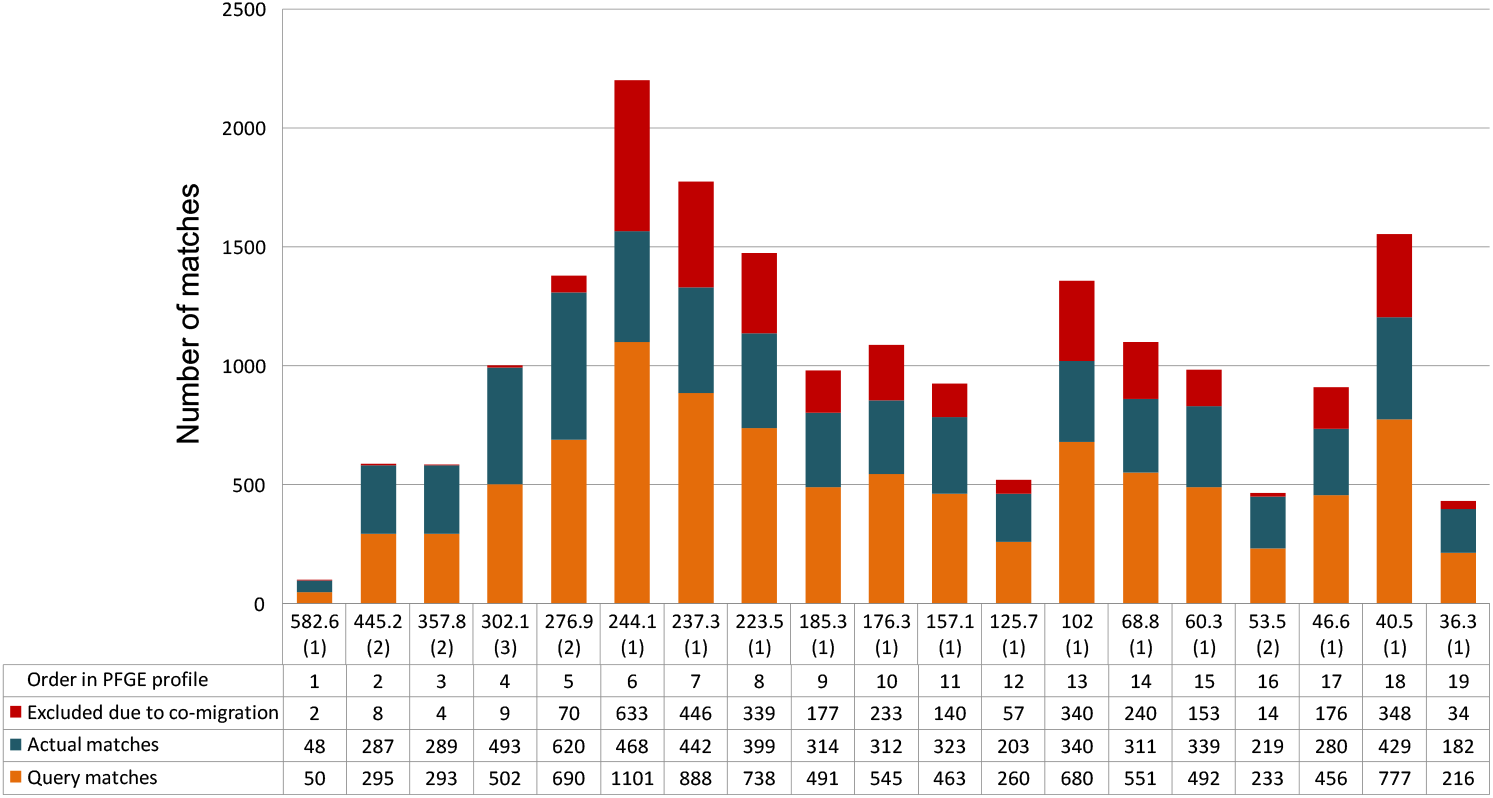
Total details of matches with each PFGE band across *Xba*I digestion profile of *E. coli* O157:H7 strain G5244 returned by GelToWGS. X-axis shows PFGE band sizes in kbp and corresponding number of DNA fragments represented by each band (FCMs).

Bands #19, #16 and #12 were also relatively rare. Interestingly, exclusion due to co-migration was very rare across PFGE bands those showed co-migration. Namely; bands #2 to #5 and #16. Single-fragment bands #1, #12 and #19 seem to be relatively rare across tested DMs. Not only because of their relatively rare occurrence, but also by their very few exclusions due to co-migrations **Fig. (1)**.

### GelToWGS comparisons with *Xba*I digestion of *S. enterica* serotype Braenderup (strain H9812)

Total query matches were 3,542 which is more than typable fragments of DMs. When considering co-migration in query fragments, the number dropped to 2,246 (74.37%). These findings suggest that co-migration reduces accuracy of PFGE by 24.64% **Table (1)**.

Among top 25 matches; matched fragments ranged from 9 to 12 while number of typable fragments per DM ranged from 23 to 29. They represented 250 fragments out of 2,246 (11.13 % from total actual matches). Dice coefficient of variation ranged from 0.5789 to 0.4615. Matches with the same serovar (Braenderup) comprised only 11 fragments (0.49 % from total). In terms of matches; top three matches belong to different serovars (two Dublin and one Typhimurium var. 5-). Match number 4 belong to the same Braenderup serovar (accession number CP022490.1) **Table (1)** and supplementary data set 2 (S2).

Epidemiological information for the previously mentioned DM (retrieved from NCBI-BioSample database Key: SAMN06044997) show that it belong to strain SA20026289. No more data is available.

When we consider matches with each PFGE band across DMs; actual matches with bands #6, #2, #9, #7 and #17 comprised 1,515 (67.54%) out of 2,246. Effect of co-migration is highest (425 out of 673) in band #6 (310.1 kbp representing a single DNA fragment) followed by #9, #7 and #2. In contrast, #16, #1, #15 and #14 have vastly less occurrences across DMs tested. Interestingly, #15 and #13 actually represent 3 and 2 different co-migrated fragments respectively. Exclusions due to co-migration for both bands were 0 and 2 respectively **Fig. (2)** And supplementary data set 2 (S2).

**Fig. (2):**
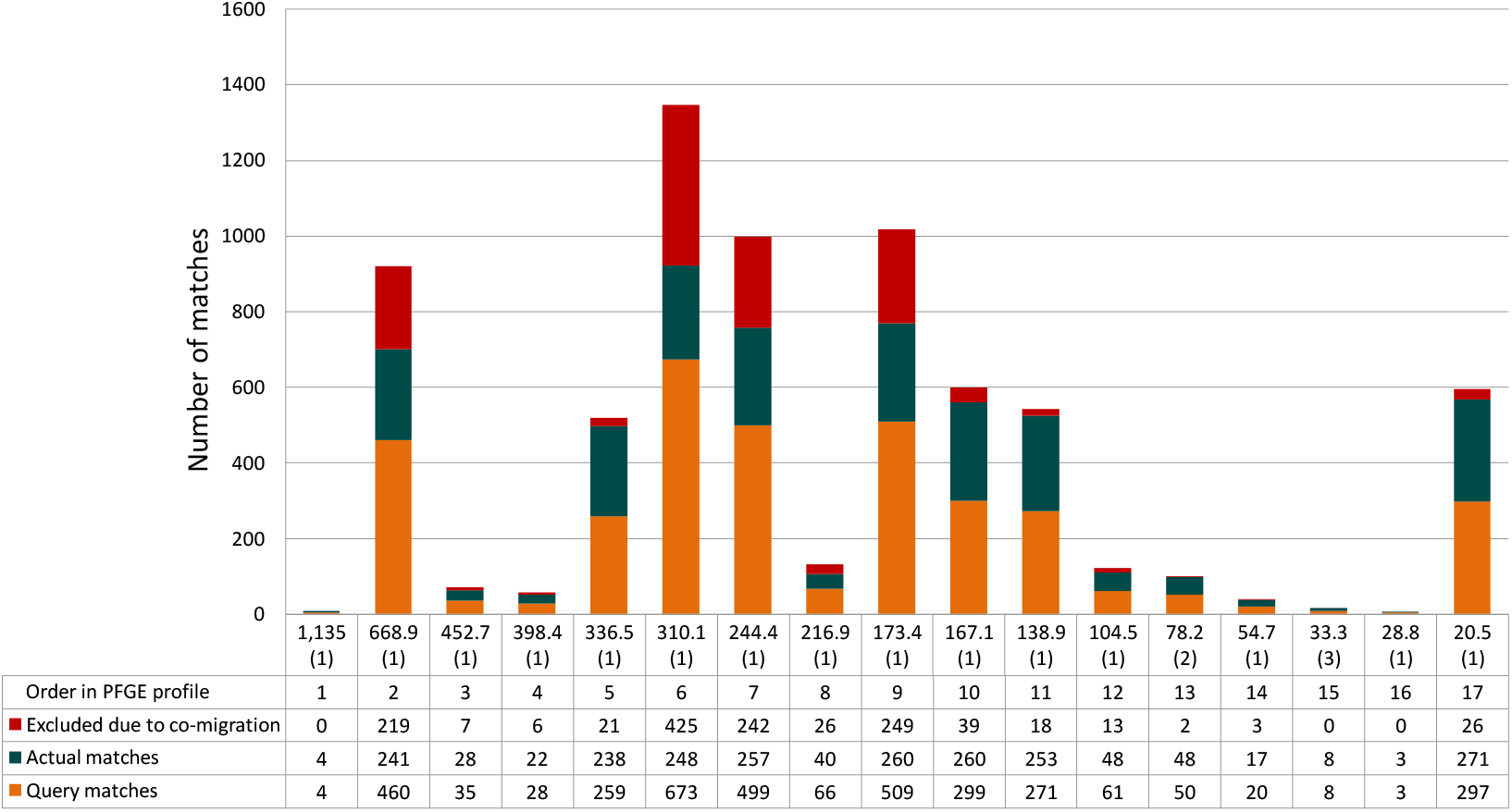
Total number of matches for each individual PFGE bands across *Xba*I digestion of *S. enterica* serotype Braenderup (strain H9812) returned by GelToWGS database. X-axis shows PFGE band sizes in kbp and corresponding number of DNA fragments represented by each band (FCMs).

Detailed comparison between test subject and the DM that belongs serovar Braenderup strain SA20026289, reveal an error in GelToWGS comparison algorithm. A single query fragment was considered a match more than once. Band #9 and #10 (173.4 and 167.1 kbp respectively) of test subject (at 3.7% CCT) had query fragment of 168.336 kbp counted twice as query band size and FCM. In spite of this error, final conclusion of actual matches is not affected, obviously because both test fragments represent a single DNA fragment according to FCM-ECSB calculations **Fig.(3)**. Interestingly, bands #1, #16, #3 and #4 which showed rare occurrences across DMs were all present in this DM which belongs to the same serovar as test subject **Fig.(3)**.

**Fig. (3):**
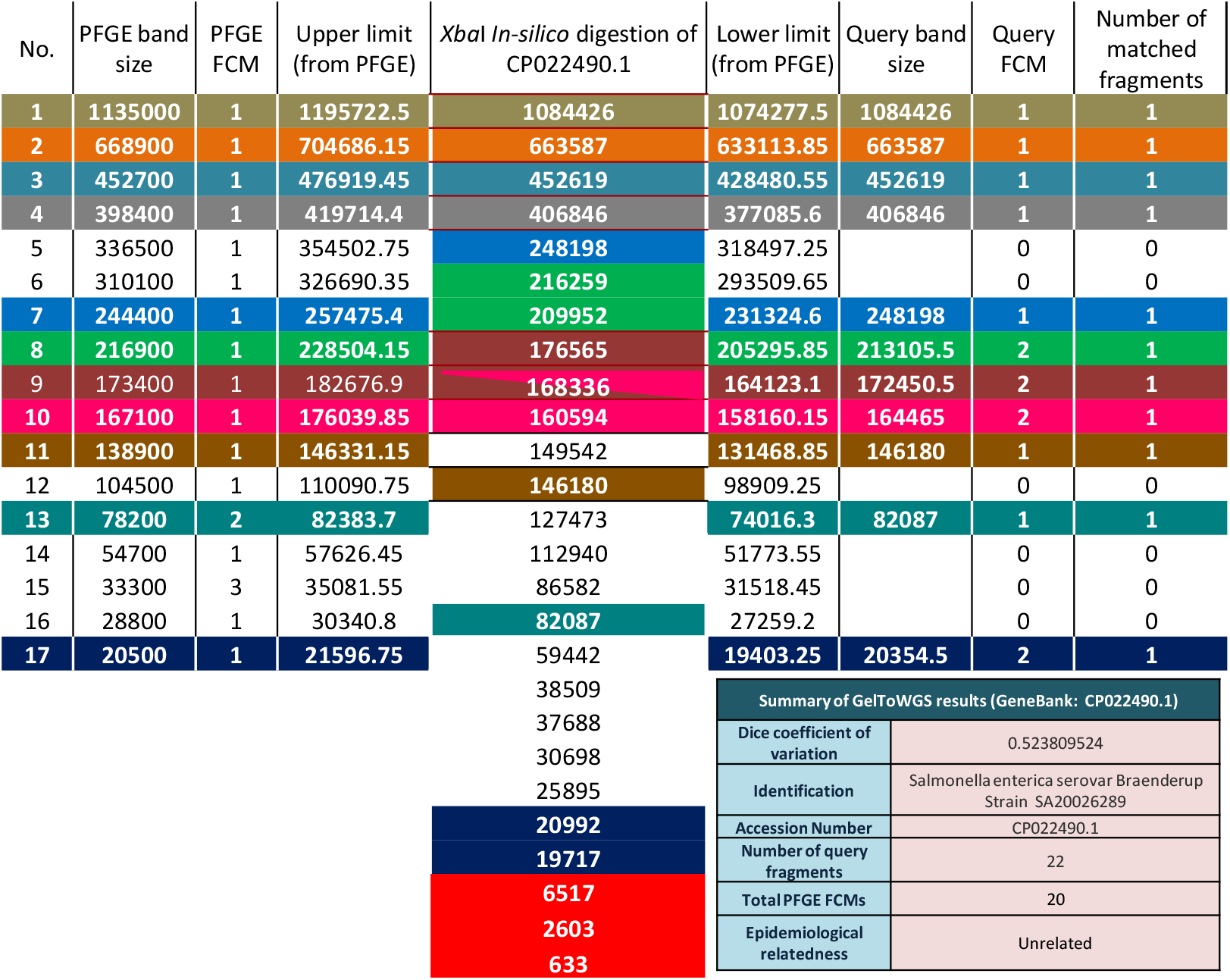
Details of comparing PFGE-FCM results of *Xba*I digestion of *S. enterica* serotype Braenderup (strain H9812) to i*n-silico* obtained *Xba*I-digested whole chromosome sequence of *S. enterica* that belong to the same serotype (strain SA20026289). NCBI-GeneBank accession number CP022490.1. Results were returned by GelToWGS database. Each match is indicated by a different color. Notice that DNA fragments of *in-silico* digestion, which corresponds PFGE band #9 considered as a match with both PFGE bands #9 and #10 (indicated by overlapping pink and dark red). Query FCMs for both corresponding PFGE bands were set to 2. DNA fragments indicated by Red backgrounds are non-typable by PFGE.

## Discussion

Since GelToWGS database is built to bridge the gap between PFGE and WGS, results shown above reveal reasonable conclusions under currently adopted criteria of PFGE evaluation [18]. A single DM of *E. coli* test was possibly related to PFGE test subject. Both of them belong to the same serovar (O157:H7). *S. enterica* test showed no relationship. We argue that number of DMs stored in the database is small; it is not representative to the diversity of these species. This claim is especially true for *S. enterica* test. A single DM within the same serovar was shown. It showed no epidemiological relationship. For *E. coli* test; a significant number of DMs within serovar O157:H7 is stored in GelToWGS database. Possibly because pathogenic forms in this species are sequenced more frequently. It is important to clarify that these conclusions were made based on DNA fragments in both; PFGE images and *In-silico* digested WGSs. Algorithmic error shown in **Fig. (3)** is obviously serious and should be corrected.

Since GelToWGS algorithms are configured to run a simulation based on the assumption that DMs were run under the same wet-lab PFGE conditions, reasonability of results would never be achieved without considering co-migration in both PFGE and DMs. It is important to deeper evaluate FCM-ECSB algorithm in terms of sequencing PFGE bands after DNA extraction from gel. If such test is done for an isolate that already been sequenced, it would be more informative. Alongside, if GelToWGS database is upgraded to store sequences of each fragment (instead of length only), PFGE typing will be upgraded to a new level. Such steps will allow epidemiologists to evaluate PFGE and *In-silico* digested WGS in terms of qualitative approach; matches may be linked to chromosomal loci of the fragment. Common fragments within serotypes and pulsotypes may become determinants of these categories **Fig. (2)**. Rare matches coupled with high overall number of matches will indicate degrees of relatedness described by Tenover and his team [18] **Fig. (1 and 2)**.

Regarding our suggested method of selecting CCT, the question raised by authors of GelToWGS still needs an answer which is “does Dice percentage of differences reflects behavior of DNA fragments during PFGE?” The answer is crucial not only to described method in this article, but also to WGSToGel simulation algorithms upon which the claim that 35% of PFGE fragments co-migrate [11] is made.

When we consider findings of WGSToGel simulations reported by the same publication (statistics from supplementary material 4) of the cited article [11]. *Xba*I DMs of *E. coli* and *S. enterica* (64 and 70 DM respectively) suggest that recognition sequence of the enzyme (…TCTAGA…) [20] occurs across *E. coli* chromosomes more than *S. enterica* **Table (2)**. Consequently, co-migration occurs more frequently in *E. coli* DMs. When considering limitations of PFGE; averages and standard deviations clearly reflect effects of marker band size limitation and co-migrations (total number of fragments compared to both; PFGE- typable and expected bands) **Table (2)**. Considering FCM-ECSB results of test subjects, sum of FCM represents typable fragments, while number of bands correspond expected ones. FCM-ECSB results are within the ranges of maximum and minimum values of WGSToGel simulations; 19 bands representing 25 fragments for *E. coli* strain G5244 and 17 bands showing 20 fragments in *S. enterica* strain H9812 **Table (2)**.

**Table (2):** Summary statistics of WGSToGel simulations of *Xba*I digestions of whole chromosome sequences of *E. coli* and *S. enterica*.

| PFGE simulation | Statistics | %age of typable size | Total Number Of Fragments | PFGE-typable Fragments | Expected PFGE bands |
| --- | --- | --- | --- | --- | --- |
| *XbaI* digestion model of *E. coli* whole chromosome sequences (64 in total) | Sum | N/A | 3609 | 2419 | 1379 |
|  | Mean<br>± SD | 97.3045<br>± 3.4307 | 56<br>± 41 | 38<br>± 18 | 21<br>± 5 |
|  | Max | 99.692 | 161 | 82 | 32 |
|  | Min | 82.451 | 17 | 4 | 4 |
| *XbaI* digestion model of *S. enterica* whole chromosome sequences (70 in total) | Sum | N/A | 3412 | 2426 | 1551 |
|  | Mean<br>± SD | 95.498<br>± 9 | 49<br>± 21 | 35<br>± 12 | 22<br>± 6 |
|  | Max | 99.845 | 73 | 63 | 42 |
|  | Min | 66.694 | 18 | 16 | 12 |

Data in this table were derived from a supplementary material 4 (S4) of previously published article by IEM Adam and his team. The article described WGSToGel simulations of several PFGE protocols.

